# Multidrug Efflux in Gram-Negative Bacteria: Rationally Modifying Compounds to Avoid Efflux Pumps

**DOI:** 10.1101/2023.07.13.548850

**Authors:** Dominik Gurvic, Ulrich Zachariae

## Abstract

Gram-negative bacteria cause the majority of critically drug-resistant infections, necessitating the rapid development of new drugs with Gram-negative activity. However, drug design is hampered by the low permeability of the Gram-negative cell envelope and the function of drug efflux pumps, which extrude foreign molecules from the cell. A better understanding of the molecular determinants of compound recognition by efflux pumps is, therefore, essential. Here, we quantitatively analyse the activity of over 73,000 compounds across three strains of *E. coli* – the wild-type, an efflux-deficient variant, and a hyper-permeable variant – to elucidate the molecular principles of evading efflux pumps. Our results show that, alongside a range of physicochemical features, the presence or absence of specific chemical groups in the compounds substantially increases the probability of avoiding efflux. Furthermore, comparison of our findings with inward permeability data highlights the primary role of efflux in determining drug bioactivity in Gram-negative bacteria.

## Introduction

Gram-negative (GN) bacteria are responsible for the majority of highly or extremely drugresistant bacterial infections. For the most critically resistant pathogens, drug treatment strategies are now limited to few remaining options, and therefore new or improved antibacterials are urgently needed. ^1–4^ However, only a small number of antibacterial drugs are currently under development, whose activity is moreover strongly biased toward Grampositive (GP) bacteria.^5,6^ The architecture of the GN cell envelope, consisting of two lipid membranes which enclose the periplasmic space, represents the major obstacle preventing sufficient drug activity in GN bacteria.^7–9^ By contrast, GP bacteria only possess a single membrane. The chemical determinants for enhanced drug permeation across the GN cell envelope remain largely unclear, which substantially hinders the design of new drugs with GN activity.^10,11^

The two lipid membranes in the GN cell envelope, the outer membrane (OM) and cytoplasmic membrane (CM), contain key proteins which play an important role in drug permeability (see Fig. 1). Inward drug permeation across the OM is thought to proceed primarily via porin proteins, whose pores possess specific geometries and are highly polar with a strong transversal electrostatic field, limiting the chemical space available for permeant molecules.^12,13^ Porins can undergo mutations, and their expression level can be down-regulated to enhance resistance.^10,14–16^ For many GN bacteria, however, active drug efflux is thought to be a major driver of intrinsic and acquired drug resistance. ^17–19^ In particular, GN bacterial tripartite efflux pumps, which span both the CM and the OM and the periplasm, efficiently recognise and expel most drugs from the bacterial periplasm before they are able to reach their therapeutic targets - Fig. 1.^20–22^

**Figure 1:**
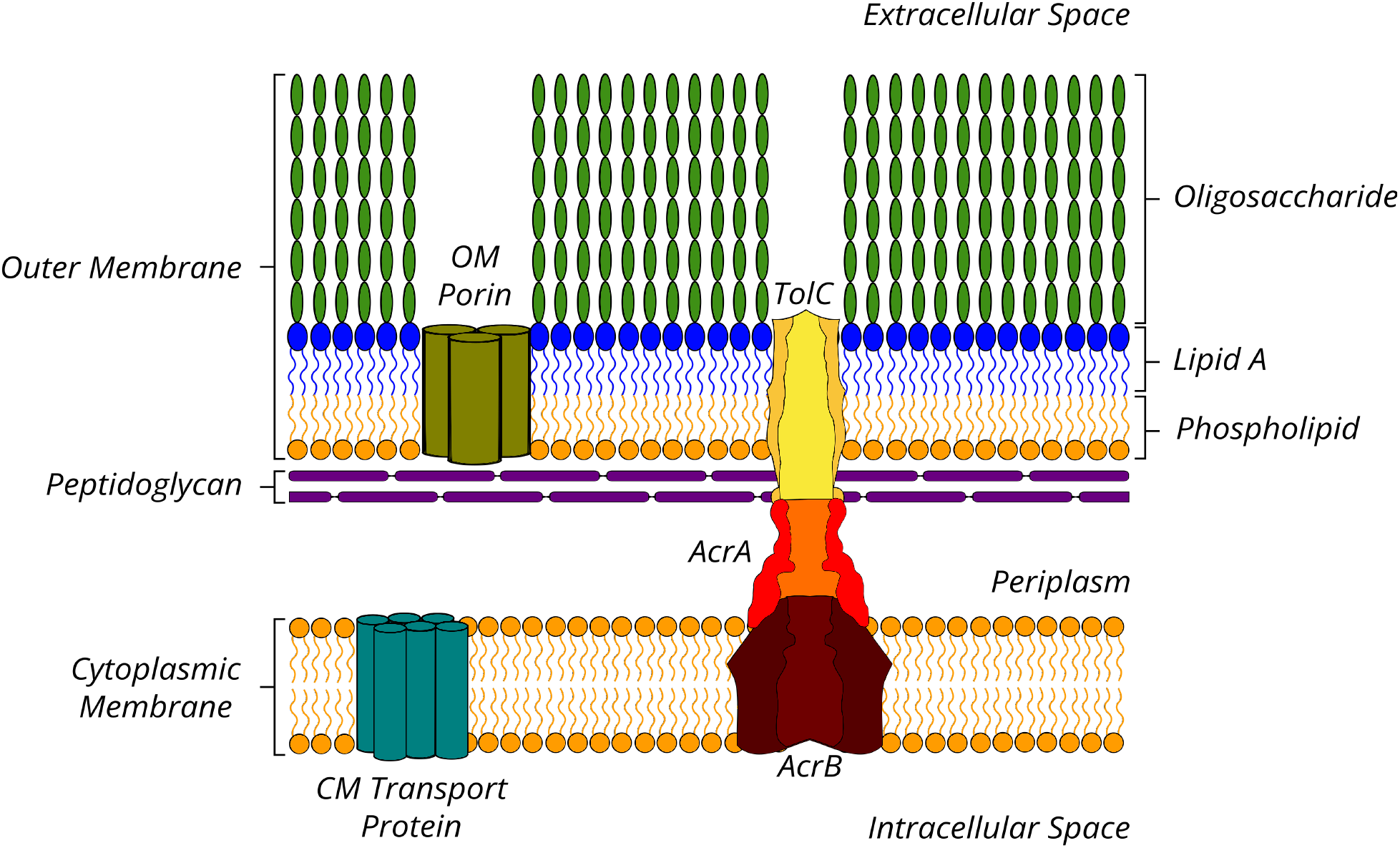
Diagram of the Gram-negative cell envelope. The outer and inner membranes enclose the periplasm with the in-lying peptidoglycan layer. Porins provide entry passageways for polar molecules into the periplasm. Tripartite efflux pumps (here: the *E. coli* AcrAB-TolC complex) span the inner and outer membranes and efficiently expel drugs from the periplasm.

Tripartite efflux pumps are driven by electrochemical gradients across the CM or by ATP hydrolysis in the cytoplasm. ^17,23^ They consist of an active pump protein in the CM, an adaptor protein in the periplasm, and an outward conduit protein in the OM, with varying stoichiometries.^21^ The spectrum of efflux substrates is notoriously broad, and a range of different pumps operate in parallel.^17,24,25^ In Enterobacteriaceae, however, including the best understood GN bacterium *Escherichia coli*, only one gene encodes an OM conduit protein, TolC.^26–28^ Consequently, all parallel efflux pathways based on tripartite efflux pumps can be inhibited by the deletion of this gene in *E. coli*.^21^

So far, the understanding of active efflux remains insufficient. It has proven especially difficult to infer specific rules that could help to guide the molecular design of efflux-avoiding compounds, which could improve the drug discovery rate for GN bacteria.^29–31^ To accelerate drug discovery in this area of considerable medical need, it is thus important to elucidate the structural and physicochemical determinants of effluxed compared to non-effluxed compounds. In this work, we therefore conducted a large-scale, data-driven chemical analysis of the experimental compound activity in different *E. coli* strains. We focused in particular on *∼*74,000 compounds acting on wild-type *E. coli* (WT; strain ATCC 25922), the efflux-deficient variant *tolC* (strain MB5747), and the OM-permeable variant (*lpxC*, strain MB4902). In *lpxC E. coli*, the lipopolysaccharide content of the otherwise poorly permeable outer leaflet of the OM is reduced, generating porin-independent permeation pathways.^32–34^

Comparing the activity of each compound across the three variants allowed us to gain insight into the balance of the two major factors underpinning low drug uptake in GN organisms, active efflux and OM permeation, and potential differences in their chemical and physical determinants. Using Matched Molecular Pair Analysis (MMPA), we identified small molecular changes that are repeatedly observed to covert a given compound from an efflux pump substrate into an efflux pump evader while at the same time not being prone to inward permeability issues. These molecular modifications and physicochemical guidelines may help medicinal chemists and drug designers to rationally enhance drug uptake in GN bacteria, which has so far represented a major obstacle to the development of GN-active antibiotics.

## Results

### Compound Classification

We sourced experimental growth inhibition (GI) data, reflecting the activity of each investigated molecule in WT as well as the *tolC* and *lpxC* variants of *E. coli*, from the public-domain CO-ADD project database (www.co-add.org).^35^ Since there can be substantial variation in assay conditions and results between different laboratories, we opted for consistent, single-source data recorded in-house at CO-ADD, which we deemed optimal for subsequent analysis. The data sets consist of GI values obtained as optical density values (*OD*_600_) after treatment of bacterial cultures with the compounds. The values are expressed as a percentage of inhibition by normalising the test compound *OD*_600_ values to those obtained for bacteria without inhibitors (0% GI) and media only (100% GI). Note that values outside the 0–100% range can occur as, for example, some compounds could enhance growth.

To classify any given compound as active, we applied a strict criterion to prevent noise by setting the threshold to a level that exceeds the mean compound activity in the tested set (*µ*) by four standard deviations (*σ*) (*GI ≥* [*µ* + 4*σ*]). We then classified compounds as efflux substrates or efflux evaders as follows: Compounds active against WT *E. coli* at the 4*σ* level that at the same time showed activity against *tolC E. coli* at the same level were classed as efflux evaders. The rationale for this classification was that, as judged from the GI data, compound activity did not depend on the function of TolC-dependent efflux mechanisms. Efflux substrates were then identified in the compound data by being active in the *tolC* strain but inactive in WT *E. coli*, such that the activity of these compounds was likely to be suppressed by TolC-dependent efflux. Accordingly, compounds inactive in both WT and *tolC E. coli* were classed as generally inactive, while compounds that were active in WT but inactive in *tolC* were classed as WT-only active. The activity thresholds and distribution of the compounds are shown in Fig. 2 and the classification scheme we used in Fig. 3. The classification of initially 73737 compounds resulted in 200 efflux evaders, 760 efflux substrates, 53 WT-only actives and 72724 inactives.

**Figure 2:**
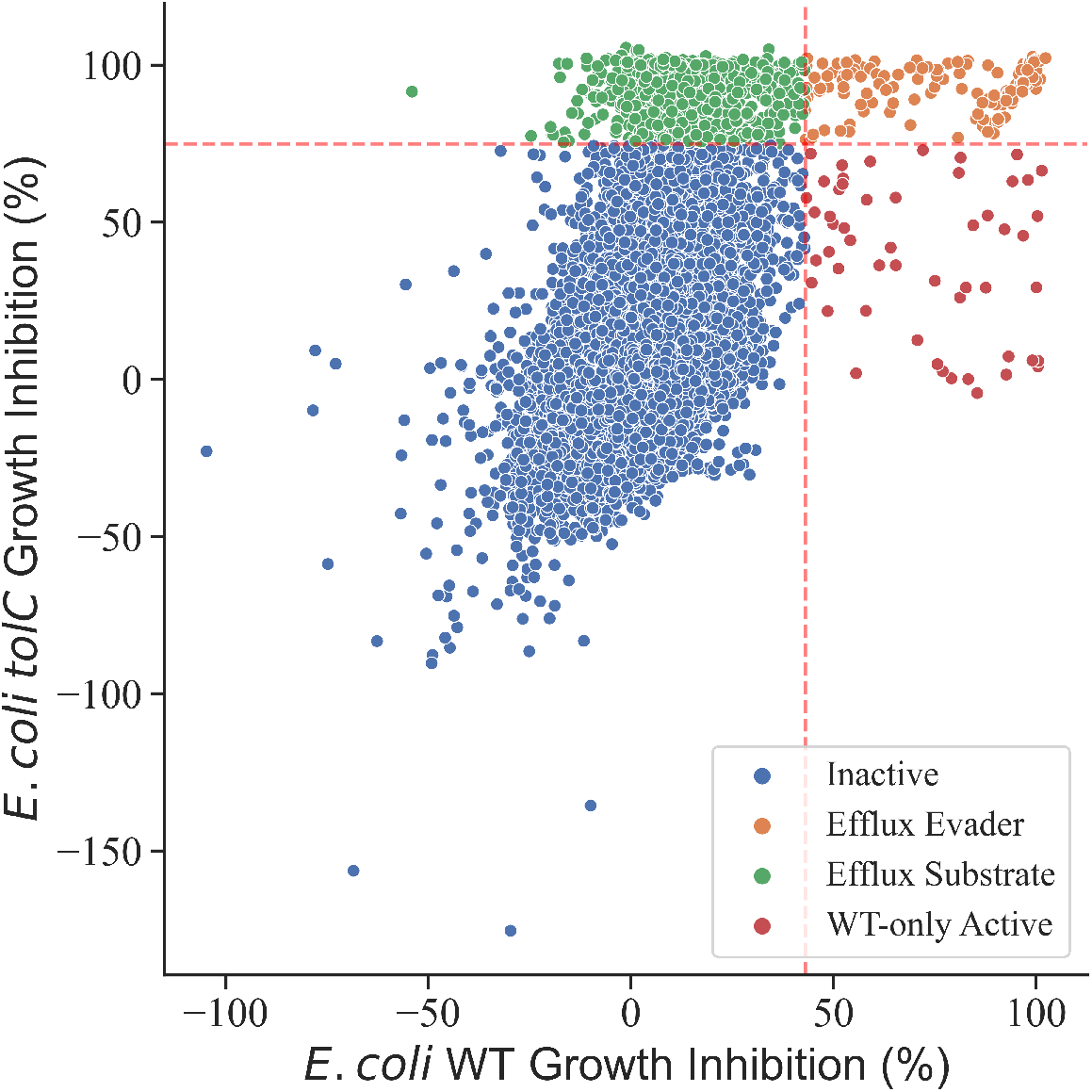
Graph displaying the experimentally determined growth inhibition values for each investigated compound in WT (x-axis) vs. *tolC E. coli* (y-axis). The thresholds for classification are shown as red dashed lines, classification results are shown in colour code. (Additional distribution of WT and *tolC*: Fig. S1)

**Figure 3:**
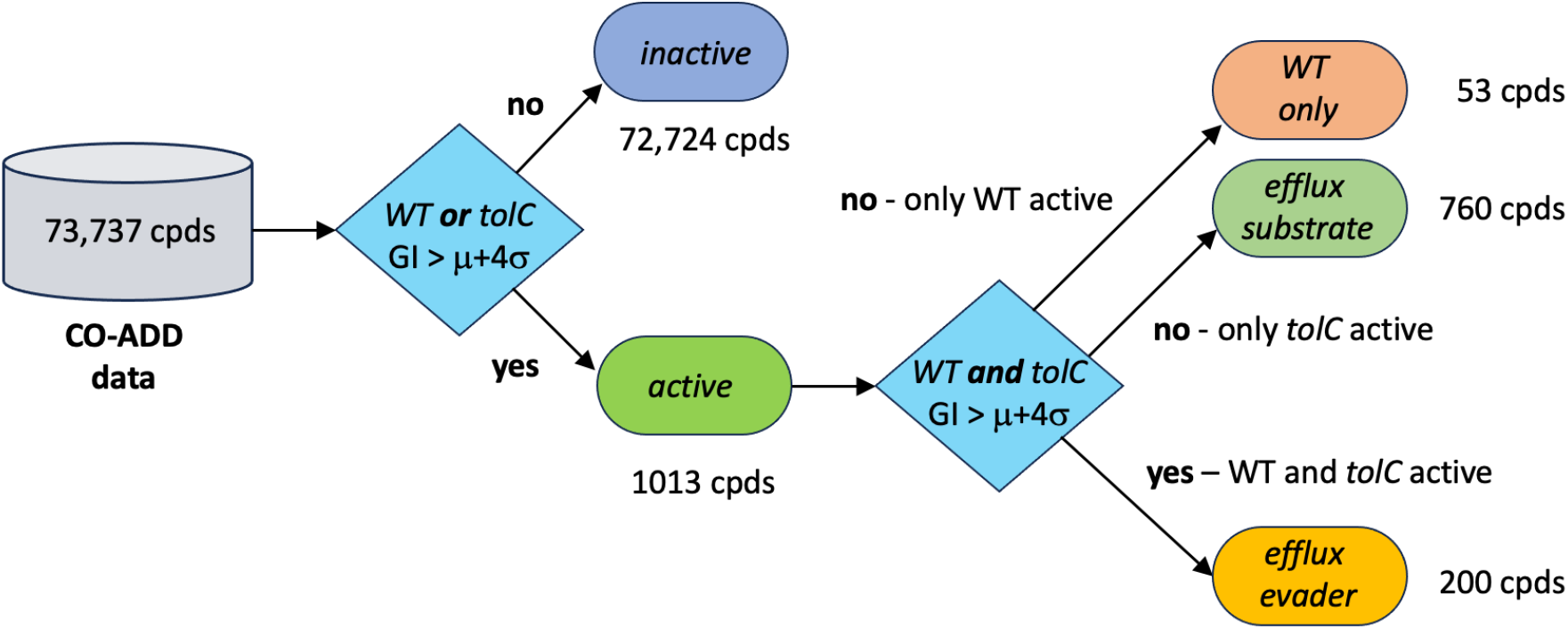
Flow chart of the compound classification procedure used to define efflux substrates and efflux evaders in addition to WT-only actives and inactives according to growth inhibition data from WT and *tolC E. coli*.

As mentioned above, it is highly challenging to identify compounds with sufficient activity against GN bacteria.^36^ The data set is, therefore, strongly biased towards the inactive class. However, the rigorous curation performed here yielded a sufficiently large data set of particularly high quality containing the most relevant compound classes for the present study, efflux substrates and evaders. The significance of the smallest class, WT-only active, is intuitively less clear. It is conceivable, however, that WT-only active compounds interact directly with parts of the efflux machinery of the WT or the tolC gene.

The data classified according to efflux characteristics then underwent a further curation step accounting for the inward permeability of the compounds across the OM. Similar to efflux, the compounds were classified into OM permeating, OM non-permeating, WT-only active and inactive molecules by evaluating activity differences between WT *E. coli* and the OM-hyperpermeable *lpxC* variant (see Supplementary Fig. S2, Fig. S3, Fig. S4). Out of the 760 compounds classified as efflux substrates, 206 were additionally found to be OM nonpermeable. For these compounds, it is difficult to ascertain whether efflux or the inability to permeate the OM is the major contributor to their WT inactivity. Hence, these 206 compounds were excluded from further analysis.

Similarly, out of the 200 efflux evaders, 186 were also classified as OM permeators, while the other 14 were found to be WT-only active with respect to OM permeation. Although, again, it is challenging to rationalise activity only in the WT, these 14 compounds were not investigated further to avoid any convolution of efflux with other effects. After accounting for OM bias, we, therefore, took 186 efflux evaders and 554 efflux substrates into further analysis (for a complete list of efflux evaders and substrate see Table S4).

Notably, examining the relative impact of low OM permeability vs. active efflux on the bioactivity of compounds, we found that a compound’s inactivity in WT can be explained by insufficient inward permeation across the OM in 369 cases, while it can be linked to active efflux in 760 cases. Hence, according to the data set investigated here, active efflux contributes to low GN bioactivity at a ratio of 2:1, as compared to overcoming the OM barrier. This shows that, indeed, active efflux is the predominant factor in determining low drug uptake across the GN cell envelope.

### Collective Physicochemical and Structural Differences Between Efflux Substrates and Evaders

We next examined if physicochemical or structural rules can be established to differentiate efflux evaders from substrates. Many efflux systems in Gram-negative bacteria recognise a notoriously promiscuous spectrum of substrates.^20,37^ Due to the existence of parallel efflux pathways and several substrate binding sites in some efflux pumps, they bind and expel drugs of a multitude of different chemotypes.^17,38^ We thus first investigated if there are potential gaps in the recognition of molecules based on their physicochemical parameters.

In earlier work, it has been suggested that compounds with whole-cell activity in GN bacteria are more hydrophilic, often zwitterionic, smaller, and have a larger polar surface area than their GN-inactive counterparts.^18,39–44^ Recent studies have also concluded that compound rigidity, represented by a number of rotatable bonds below five in the molecule, aids accumulation in GN bacteria.^36,45^ Additionally, structural elements such as various amines, thiophenes, and halides have been linked with increased GN bacterial permeation.^45,46^

In many previous studies, the shift towards increased hydrophilicity and rigidity, as well as lower MW, has been explained by the geometric and physicochemical constraints imposed by OM permeation via porins.^12,45^ For efflux in particular, recent investigations have also suggested that hydrophobic compounds are more likely to be actively expelled, whereas small hydrophilic or charged compounds and polar zwitterions have a higher probability of avoiding efflux.^39^ However, the authors note that “simply designing polar compounds was not sufficient for antibacterial activity and pointed to a lack of understanding of complex and specific bacterial penetration mechanisms”.^39^ Recently, additional features contributing to the efflux susceptibility of compounds in *E. coli* have been identified, including planarity and a greater degree of elongation with limited branching.^40^

Based on the large dataset investigated in the present study, we calculated physicochemical descriptors of the molecules for efflux substrates, efflux evaders, and a sample set of 500 inactive molecules for comparison. They include molecular weight (MW), hydrophobicity (*logP* and *logD* _7.4_), topological polar surface area (TPSA), solubility (*logS*), the number of hydrogen bond acceptors and donors, and the number of rotatable bonds (Fig. 4A). We found that the efflux evaders in our investigated data set, indeed, tend to be more hydrophilic (*logP* and *logD*), possess a larger polar surface area, and have greater solubility in water. They also have, on average, a slightly lower MW, whereas only a small difference is observed in their flexibility as compared to efflux pump substrates. However, the distributions of these features for the two classes show significant overlaps in most cases, making them poorly separable (Fig. 4A, Supplementary Fig. S6). The best separation is seen for *logP* and *logD*, where nearly no efflux substrates are observed with a *logP* below 1 or a *logD* below 0, that is, the probability of evading efflux is substantially increased for very hydrophilic compounds.

**Figure 4:**
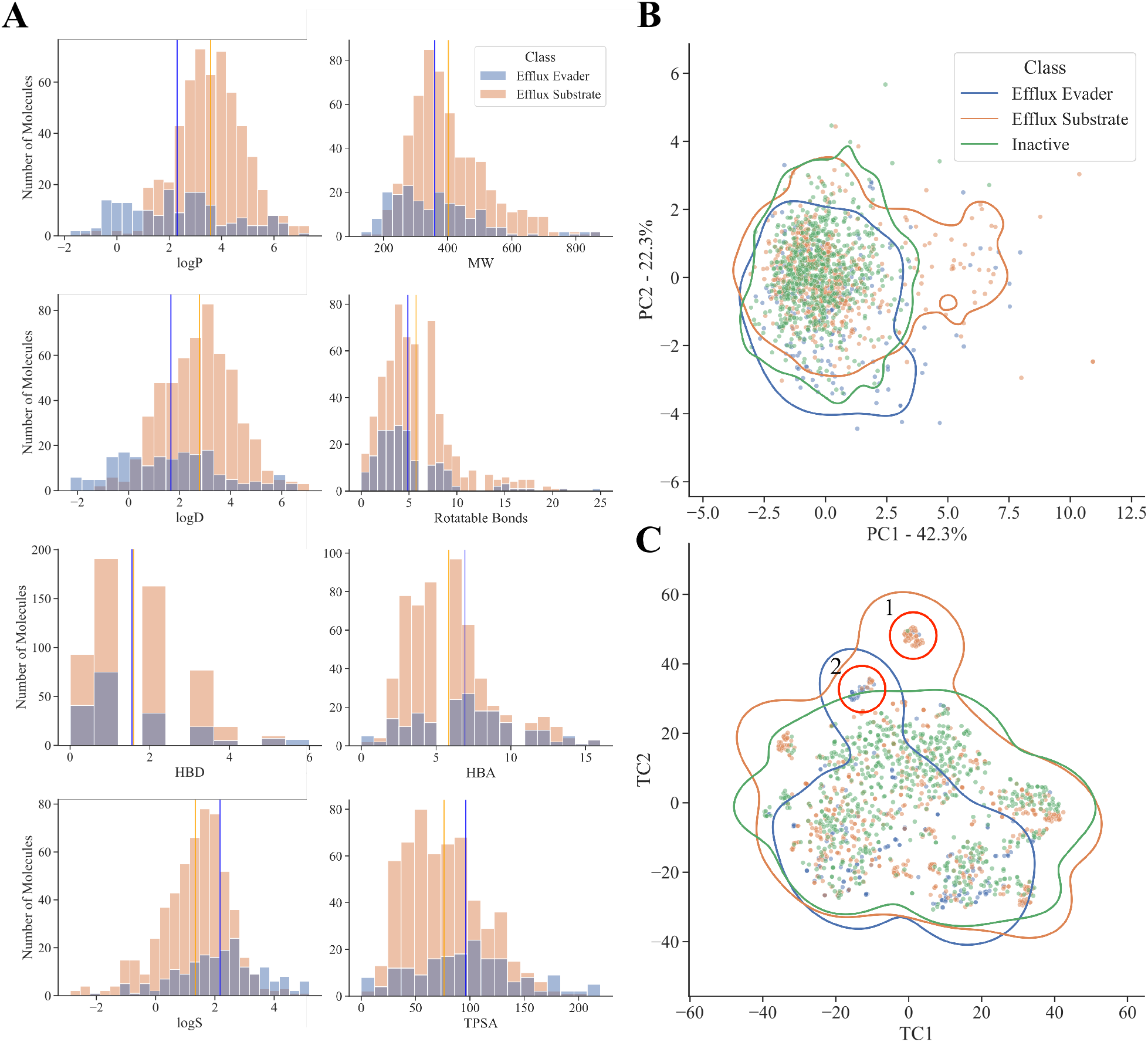
(A) Distribution of eight key physicochemical features for efflux evaders (blue) vs. efflux substrates (orange). The mean values of each distribution are shown as vertical lines. (B) Principal component analysis using the same set of physicochemical features for efflux evaders, substrates and inactive compounds. The first two principal components cover 64.6 % of the variance. (C) Comparison of the chemical similarity space occupied by efflux evaders, substrates and a sample of inactive molecules by t-SNE analysis. Two distinct clusters are marked by red circles. The contour lines in (B) and (C) were determined using a kernel density estimator and drawn at a density value of 0.05.

A principal component analysis (PCA) of the entire set of calculated physical and chemical molecular features, by contrast, showed that efflux evaders and substrates occupy largely the same space. The largest contributor to principal component 1 (PC1) was the topological polar surface area (TPSA), while for principal component 2 (PC2), it was compound hydrophobicity (*logP*; Fig. 4B). Importantly, a sample of 500 inactive compounds also displayed a substantial overlap with efflux evaders and substrates in the physicochemical feature space. This indicates that compound activity and interactions with efflux pumps may not be sufficiently explainable by a combination of collective physicochemical features alone and that, additionally, molecular structural information must be taken into account.

We, therefore, next used t-distributed Stochastic Neighbour Embedding (t-SNE) to reduce the dimensionality of the compound structural data to a two-dimensional representation according to their structural similarity (Fig. 4C). The t-SNE plot shows that efflux evaders, substrates, and inactives again occupy largely the same similarity space, demonstrating that structurally similar molecules interact differently with the efflux pumps. Of note, however, are two small separated clusters of structurally similar molecules at the top of the t-SNE plot (Fig. 4C, Clusters 1 and 2). Cluster 1 consists of 40 compounds, three of which are efflux evaders, while 37 are efflux substrates. Cluster 2 displays a ratio of 23 efflux evaders vs. 9 efflux substrates, none of which possess similarity to any of the 500 sampled inactive molecules (the structures belonging to the two clusters are shown in Table S2, the maximum common substructures characterising each cluster in Fig. S7). A large proportion of the compounds in cluster 2 contain a carboxylic acid moiety. However, we were not able to identify any molecular transformations within this cluster that convert an efflux substrate into an efflux evader by adding a carboxylic acid group (for details on molecular transformations, see below).

In summary, we conclude that it is – similar to the observations we made for the physicochemical parameters – challenging to predict the interaction of the three compound classes with efflux pumps solely on the basis of collective molecular features such as structural similarity. We, therefore, next investigated if more fine-grained differences on the level of small structural substitutions may be decisive for the recognition of compounds by efflux pumps.

### Molecular Transformation of Compounds Between the Inactive, Substrate and Evader Classes

We used matched molecular pair analysis (MMPA) to analyse small structural changes that convert compounds with a common core between the three compound classes; inactives, efflux evaders, and efflux substrates (Fig. 5). The analysis yielded a set of 4900 substrate transforms, in which 2053 inactive compounds are transformed into 349 substrates, as well as a set of 612 transforms in which 397 inactive compounds are converted into 77 efflux evaders. Note that in all cases, the number of transforms exceeded the total number of the compounds since, in many cases, multiple compounds are transformed into the same substrate or evader, while individual compounds can also be transformed into multiple substrates or evaders (see Table 1). Overall, due to the smaller number of efflux evaders in the initial data set, there are fewer transforms leading to evaders than to substrates, while the number of inactive compounds exceeds both substrates and evaders.

**Figure 5:**
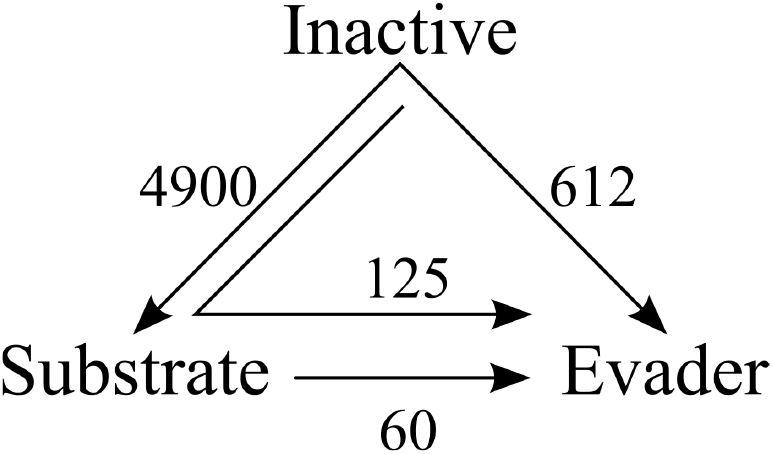
Molecular transformations resulting from Matched Molecular Pair Analysis (MMPA). The transformations connect compound pairs converting similar compound cores from inactive to efflux substrate, inactive to efflux evader, and from substrate to evader (arrows). The numbers of independent transforms are shown next to each arrow. The arrow inside indicates a double transform, from inactive to substrate and further to an evader, connecting compound triplets.

**Table 1:**
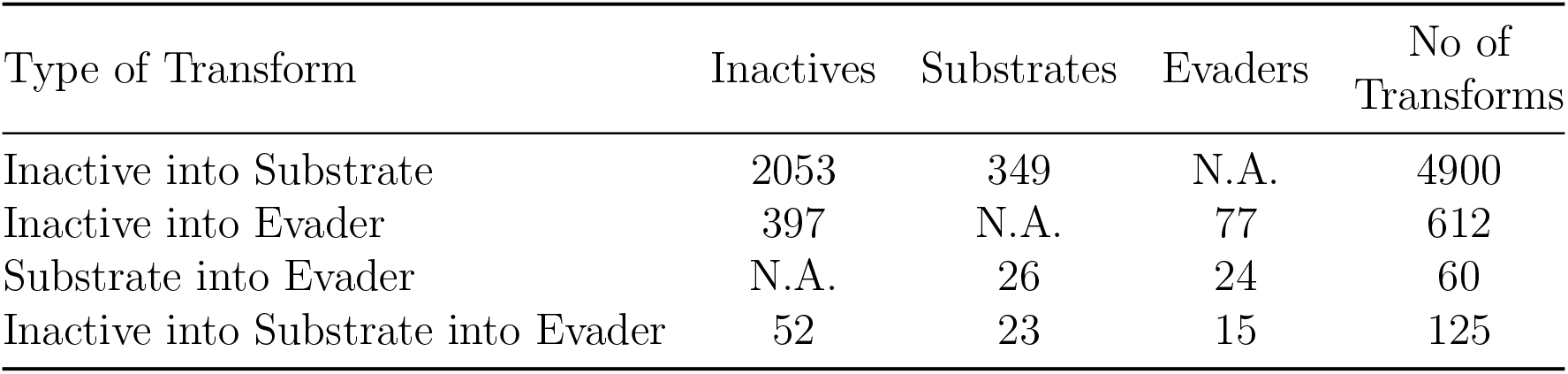
All transformations from MMPA, together with the number of unique compounds from each class within the transforms. Multiple transforms can occur between similar compounds, such that the number of transforms exceeds the number of classified compounds.

Central to the main focus of our study, 60 of the identified transforms converted 26 substrates into 24 evaders. Moreover, 125 double transforms linked 52 unique inactive compounds to 23 substrates and, further, to 15 evaders by consecutive molecular substitutions, connecting compound triplets rather than pairs. Each individual sequence of transforms in these 125 examples contained an identical core and three replacement moieties attached to the same location on the core, resulting in members of the three different compound classes.

Fig. 6 shows eight exemplar double transforms (all transforms are provided in the SI - Table S3). The associated WT and *tolC* activity values highlight the large impact of small substructural changes on the compounds’ bioactivity and their interaction with efflux pumps.

**Figure 6:**
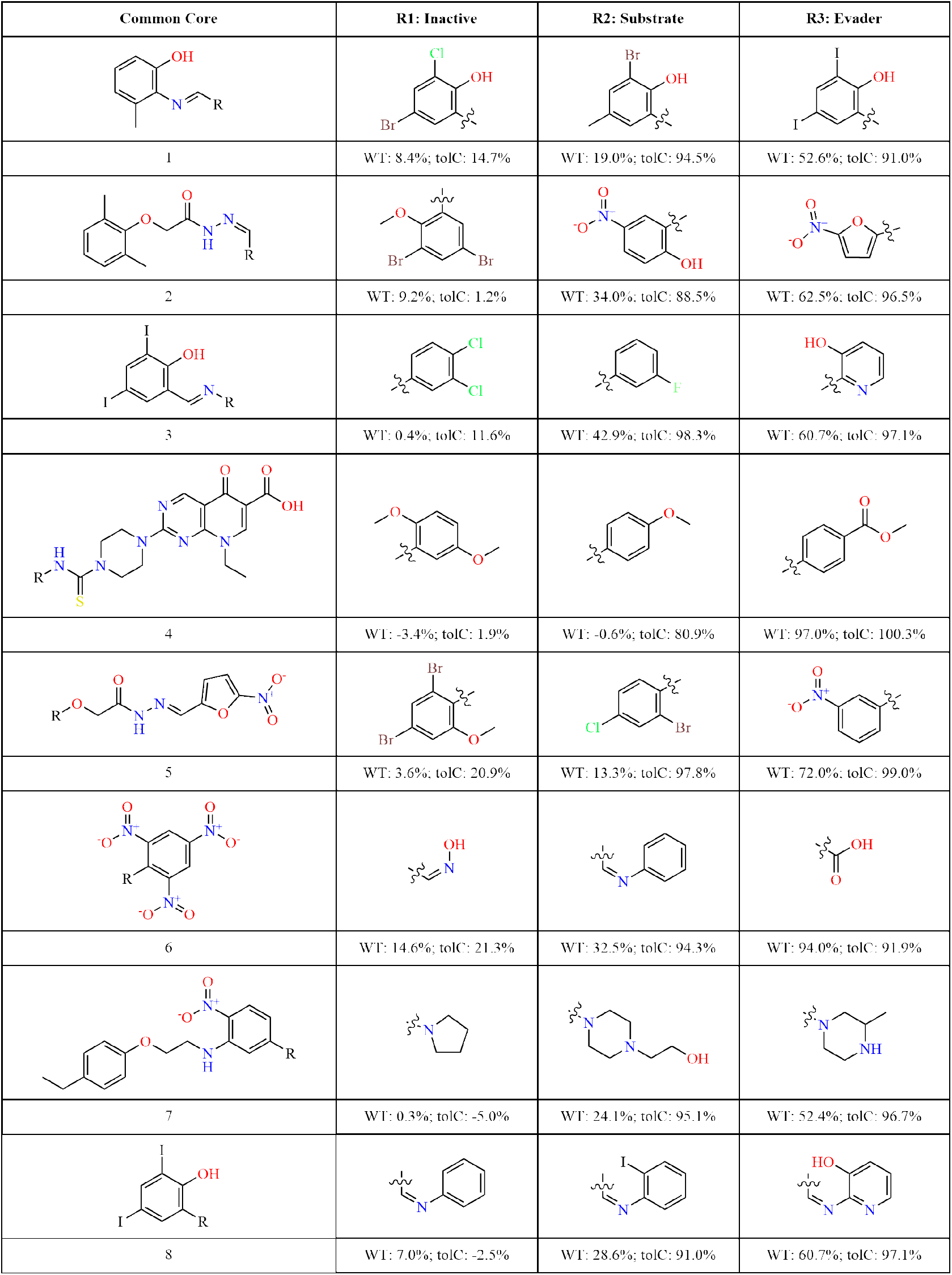
Exemplar transformations converting inactive molecules into efflux substrates and further into efflux evaders by small structural modifications between these compound triplets. The first column shows the compound core shared by all three molecules in the triplet, and the following three columns show the substitutions resulting in inactive, substrate and evader compounds. ‘R’ on the common core structure indicates the location of the molecular substitutions in column 1; ‘R1’ is the substitution required for inactives, ‘R2’ for substrates, and ‘R3’ for efflux evaders. Each compound is labelled with its associated GI values measured for WT *E. coli* and the *tolC* variant, respectively.

### Transforming Efflux Substrates into Efflux Evaders

Focusing on molecular transforms that turn efflux substrates into evaders, we searched for recurring patterns in which addition or removal of chemical moieties led to this conversion (Table 2). We found that within the transforms, the addition of pyridine was seen to transform efflux substrates into efflux evaders in 22 individual cases. Likewise, adding primary, secondary, or tertiary amines (either aliphatic or aromatic amines) converted substrates into evaders in 16 independent transformations. In addition to these nitrogen-containing groups, adding *α*-halogenated carbonyl groups was found to turn substrates into evaders in four cases. Conversely, looking at chemical moieties whose removal promoted efflux evasion, in addition to the more exotic iodo-group, we found aromatic alcohols (10 repeats), quaternary ammonium cations (7 repeats), ketone and aldehyde groups (altogether 6 repeats), as well as ether groups (4 repeats). These groups, therefore, enhanced a compound’s recognition as an efflux substrate.

**Table 2:** Moieties whose addition to or removal from a common core promotes a compound’s avoidance of efflux. The moieties have a positive effect on efflux evasion when their addition results in an evader and a negative effect when their removal results in an evader. The number of transforms in which the same moiety and effect are observed is shown as repeats.

| Moieties | Effect | Repeats |
| --- | --- | --- |
| Pyridine | Positive | 22 |
| 1°/2°/3° aromatic and aliphatic amine | Positive | 16 |
| Alpha-halogenated carbonyl | Positive | 4 |
| Iodo-moieties | Negative | 11 |
| Aromatic alcohols | Negative | 10 |
| Quaternary ammonium | Negative | 7 |
| Alkyl and aryl ketones and aldehydes | Negative | 6 |
| Ether | Negative | 4 |

The overall picture that emerges from transforming efflux substrates into evaders is that replacement of oxygen-containing functional groups with nitrogen-containing groups aids in efflux evasion, with the exceptions of quaternary ammonium cations (negative correlation with efflux evasion) and *α*-halogenated carbonyls (positive correlation with efflux evasion). This effect is likely due to specific molecular features of the substrate binding sites in the bacterial efflux pumps.

Notably, the largest contributor to the transformation of compounds from inactives to both substrates and evaders is the nitro group. The polar nitro moiety appears in 62 evader compounds and in 138 substrate compounds, such that the addition of the nitro moiety to an inactive compound can lead to both evaders and substrates. It is noteworthy, however, that a remarkable enrichment of evaders is observed amongst the nitro-containing active compounds, occurring in 62 out of 186 analysed evaders (33%).

To ascertain if the moiety exchanges identified within the transformations conformed to the general shifts in the physicochemical parameters observed for efflux evaders vs. substrates (see Fig. 4), we examined their average change amongst all transformations. (Table 3) shows that the evaders were smaller than their substrate counterparts, with an average decrease of around 37 Da (−10.7%) in the transformations. Evaders were also more hydrophilic than substrates, with a *logP* and *logD* decrease of −33.3% and −38.1%, respectively. On average, the number of rotatable bonds decreased, albeit by a small percentage (−5.8%), indicating that evaders tend to be more rigid. The solubility, *logS*, increased (15.5%) in line with a lowered *logP*. A raised TPSA (21.8%) indicates a gain in potential polar interactions, again in agreement with a decreased *logP*. The largest magnitude increase was seen for the number of hydrogen bond donors (28.7%) in addition to a smaller increase in hydrogen bond acceptors (18.0%). This translates into an increase of nearly one additional hydrogen bond acceptor in evaders, on average, as compared to substrates.

**Table 3:** Average shift in physicochemical features in the molecular transforms converting compounds from substrates to evaders (change), as compared to the shift of physicochemical features in all compounds classed as substrates and evaders (bulk change) and the shifts observed for all OM non-permeable *vs.* OM-permeable compounds.

| PC feature | Substrates | Evaders | Change | Bulk Change | OM Permeation |
| --- | --- | --- | --- | --- | --- |
| MW | 348.93 | 311.53 | -10.7% | -10.95% | +0.18% |
| *logP* | 2.58 | 1.72 | -33.3% | -36.6% | -13.24% |
| *logD* | 2.26 | 1.40 | -38.1% | -40.79% | -21.15% |
| Rot. bonds | 3.24 | 3.05 | -5.8% | -15.2% | -8.81% |
| *logS* | 2.45 | 2.83 | +15.5% | +63.9% | +7.62% |
| TPSA | 71.72 | 87.37 | +21.8% | +27.14% | +1.59% |
| HBA | 5.05 | 5.96 | +18.0% | +19.14% | +5.64% |
| HBD | 0.94 | 1.21 | +28.7% | -1.72% | -3.96% |

The specific molecular transformations converting efflux substrates into evaders reproduced the bulk shifts in physicochemical parameters observed for all substrates vs. evaders (Fig. 4, Table 3), with the exception of the number of hydrogen bond donors, where a marked increase was seen for the transform pairs. Overall, the major changes in these markers point to increased hydrophilicity and polarity as key to enhancing the probability of evading efflux pumps. Taken together, these findings lead to an important conclusion: While a change in collective parameters, such as an increase in hydrophilicity, can be accomplished in various different ways by substituting hydrophobic moieties with polar moieties, our analysis shows that the *type* of hydrophilic group is the key difference between obtaining an efflux pump substrate or an efflux pump evader. More specifically, with the exception of quaternary ammonium cations, the addition of nitrogen-containing groups aids efflux evasion, whereas it is the *removal* of polar oxygen-containing moieties that yields the same effect, despite an overall average gain of hydrophilicity.

A further key insight was that, by comparing the set of physicochemical changes of efflux substrates *vs.* evaders with the physicochemical changes of OM non-permeating *vs.* OM-permeating compounds (derived from GI values measured in WT and *lpxC E. coli*), we found that the major collective parameters associated with improved GN bioactivity showed a much closer link to efflux evasion than to inward permeability across the OM. This result further supports the notion that efflux is the main contributor determining whole-cell bioactivity in GN bacteria, and that evading efflux pumps is thus key to the design of GN-active antibacterials.

## Discussion

Although drug resistance occurs in both GP and GN bacteria, GN bacteria have a much higher propensity for displaying critical or extreme resistance.^1–4^ GN bacterial infections are more difficult to treat, and drug development against GN bacteria is faced with substantial hurdles, the major one being insufficient drug uptake across the GN cell envelope.^10,11^

In the present study, we focused on the role of active drug efflux in low drug uptake. GN efflux systems expel compounds of a wide range of different chemotypes. The determinants of compound recognition by efflux pumps and, especially, their avoidance, are only sparsely understood.^17,24,29–31^ We, therefore, investigated what can be learned from the differential activity of 73,737 compounds from the CO-ADD database in three different variants of the GN bacterium *E. coli*; the WT, the efflux-deficient *tolC* strain, and the OM hyper-permeable *lpxC* strain. Many early studies of drug permeation and efflux in GN bacteria were based on smaller numbers of compounds, which additionally often showed considerable similarity to each other.^39,47^ In recent years, however, large compound activity databases such as CO-ADD have become publicly accessible, enabling the analysis of big data sets with greater statistical power.^35^

Our analysis shows that the physicochemical features commonly associated with increased general GN bioactivity of a compound mainly increase the probability of avoiding active efflux. These features include enhanced hydrophilicity, a larger polar surface area, high solubility in water, and increased H-bonding potential.^39–41,44^ In many previous studies, the increased bioactivity of these compounds had been rationalised by assuming that polar molecules are more readily able to pass through the highly charged interior of porin channels in the OM.^12,16,36,45^ In the data set we analysed, active efflux is clearly the major contributor to low GN bioactivity. The physicochemical characteristics underpinning GN activity serve to promote the evasion of efflux pumps. Comparing the relative importance of efflux vs. OM permeability in reduced WT bioactivity, about two-thirds of the compounds in our dataset active in either the efflux-deficient *tolC* strain or the OM hyper-permeable *lpxC* strain but not the WT of *E. coli* are effluxed, whereas one-third are poorly permeable across the OM. Moreover, analysis of data from the *lpxC* strain revealed that enhanced OM permeability is much less correlated with changes in physicochemical features such as polar surface area or hydrophilicity than general GN bioactivity or efflux.

Since it had been noted previously that the design of polar compounds is not sufficient for antibacterial activity,^39^ we furthermore investigated distinct chemical transformations that aid in efflux evasion. A more complex picture of efflux avoidance emerged upon a detailed structural analysis of compound pairs, in which small chemical modulations transform efflux substrate compounds into efflux evaders. Specifically, avoiding recognition by efflux pumps is linked to certain chemical modifications that conform to the physicochemical guidelines noted above, but not to alternative molecular substitutions that may have the same effect on the physicochemical parameters. For example, adding amine groups and/or pyridine, and in this way increasing hydrophilicity, is linked with the majority of molecular transformations that convert efflux substrates into evaders. By contrast, it is necessary to remove quaternary ammonium cations to achieve a similar effect. Likewise, hydrophilic oxygen-containing groups, including ketones, aldehydes, ethers, and aromatic alcohols, promote the recognition of compounds by efflux pumps.

Overall, our analysis leads to the following main conclusions: With regard to uptake through the GN cell envelope, the bioactivity of a compound in the GN bacterium *E. coli* is mainly driven by its propensity to be an efflux substrate. Strongly hydrophilic compounds with large polar surface areas and high solubility are much more likely to evade efflux; however, specific molecular modifications in the direction of increased hydrophilicity are required to escape efflux, while others have the opposite effect. In particular, nitrogen-containing functional groups including amines, pyridine, and the nitro moiety are connected with a higher probability of evading efflux.

Notable limitations of our study are the restriction to compounds tested in *E. coli* and its variants, as well as the inevitable bias between a large number of inactive compounds compared to a relatively small number of actives, of which only a subset is classified as efflux pump evaders. We selected a strict criterion for compound activity to obtain a curated data set with minimal noise. It is also important to note that not all efflux in *E. coli* is TolC-dependent. Future studies should be performed to address the chemical determinants of drug efflux within other GN pathogens and a broader range of variants from the WT.

## Methods

### Data Source

The CO-ADD microbial growth inhibition database contains information on *∼*100k compounds, from both academic and industry sources, screened against five bacterial pathogens, their mutants, as well as two fungal pathogens (*Escherichia coli* WT - ATCC 25922, *Escherichia coli lpxC* - MB4902, *Escherichia coli tolC* - MB5747, *Klebsiella pneumoniae* - ATCC 700603, *Acinetobacter baumannii* - ATCC 19606, *Pseudomonas aeruginosa* - ATCC 27853, *Pseudomonas aeruginosa* PΔ7 - PAO397, methicillin-resistant *Staphylococcus aureus* - ATCC 43300, and the fungal pathogens *Cryptococcus neoformans* - ATCC 208821 and *Candida albicans* - ATCC 90028). Inhibition of bacterial growth was determined by measuring light absorbance at 600 nm (*OD*_600_) after treatment with compounds. The percentage of growth inhibition (activity) was calculated for each assay, using media only as negative control and bacteria without inhibitors as a positive control, on the same plate as references. The growth inhibition database was downloaded from the CO-ADD website www.co-add.org.35

### Determining the Threshold for Activity

To analyse this dataset, we assigned an activity threshold that separates inactive compounds from active ones. Based on a combination of activity/inactivity of each compound in each of the *E. coli* strains, i.e., WT, *tolC* and *lpxC*, the following classes were assigned with respect to efflux pump interaction and outer membrane permeability. We defined the threshold for an active compound as an activity value at least four standard deviations greater than the mean activity value for all compounds (*≥* [*µ* + 4*σ*]). More specifically, the mean value of the percentage of growth inhibition after treatment with a concentration of each compound in WT *E. coli* is *µ* = 4.1, and the standard deviation is *σ* = 9.7. Hence, the compounds were classed as WT active if their activity is *≥* 43.0%.

The determination of the activity threshold in the *tolC* strain was performed as follows: The mean activity value was *µ* = 6.6 with standard deviation *σ* = 17.1, giving a threshold for active compounds in this efflux deficient strain of *≥* 75%. Lastly, for the *lpxC* strain, the activity threshold to classify compounds into OM-permeable and non-permeable was obtained via an activity mean of *µ* = 5.6 and a standard deviation of *σ* = 14.0. Hence, the compounds were classed as *lpxC* active if *≥* 61.8%. All the means, standard deviations and activity thresholds in their respective strains are summarised in Table 4.

**Table 4:** Determining activity threshold for three strains, in order to classify the compounds, where activity threshold is *≥* [*µ* + 4*σ*].

| *E. coli* strains | Mean ($\mu$) | Standard deviation ($\sigma$) | Activity threshold [%] |
| --- | --- | --- | --- |
| WT | 4.1 | 9.7 | 43.0 |
| *tolC* | 6.6 | 17.1 | 75.0 |
| *lpxC* | 5.6 | 14.0 | 61.8 |

Note that the threshold of at least four standard deviations is higher than those traditionally applied for this type of analysis of between two and three. However, this strict criterion allowed us to reduce potential noise from false positives.

### Physicochemical Properties and Principal Component Analysis

A principal component analysis (PCA) was carried out to reduce the dimensionality of the space spanned by all physicochemical descriptors and visualise potential differences between efflux evaders, efflux substrates, and inactive compounds in a reduced-complexity space.^48^ Rdkit was used for the calculation of PC features.^49^ Features were standardised to avoid issues due to scaling and used in linear dimensionality reduction by applying the PCA method in scikit-learn version 1.2.0.^50^ The first two principal components described a sufficient 64.6% of the variance. Physicochemical features contributing the most to the first two principal components were: the hydrophobicity measure (logP and logD), solubility measure (LogS), topological polar surface area, number of rotatable bonds, molecular weight, hydrogen bond acceptors, and hydrogen bond donors.

### t-SNE and molecular similarity

To perform the t-SNE analysis, we applied the implementation of t-Distributed Stochastic Neighbour Embedding in scikit-learn.^51^ Initially, rdkit was employed to compute Morgan fingerprints for each molecule, using a radius of 2 and generating 2048-bit fingerprint vectors.^49,52^ Subsequently, we performed t-SNE analysis with the Jaccard distance metric to reduce the data points from 2048 dimensions to two dimensions for visualisation. ^53^ The Jaccard distance is also referred to as the Tanimoto distance, and it is defined as *Tanimoto distance = 1 - Tanimoto similarity*. Therefore, the proximity between points in the t-SNE plots reflects the Tanimoto similarity of the corresponding molecules, with greater distances indicating lower Tanimoto similarity. We explored the perplexity parameter, which defines the number of nearest neighbours considered in the calculations and settled on a value of 50; the remaining parameters were used at their default values. ^50,54^

### Matched Molecular Pair Analysis

To identify small molecular differences between compounds with and without efflux pump interactions, we carried out a matched molecular pair analysis (MMPA). Matched molecular pairs (MMPs) were generated by adapting the method from Dalke et al.^55^ The results from MMPA yield pairs of similar compounds (compound A and compound B) and a small structural change between those compounds (transform), while most of the molecule (core) remains the same (Table 5). ^56^ Compound A contains the so-called left-hand-side (LHS) of the transformation, the moiety attached to the core that is replaced, while compound B contains the ‘new’ right-hand-side (RHS) chemical moiety.

**Table 5:** A single MMPA transform consists of: Compound A, built from a core and the left-hand-side (LHS); the transform then refers to changing the LHS to a right-hand-side (RHS) resulting in Compound B, which maintains the same core as Compound A, but with a new RHS attached.

| Anatomy of a transform |  |  |
| --- | --- | --- |
| compound_A | Transform | compound_B |
| core + LHS | LHS → RHS | core + RHS |

We curated all matched molecular pairs such that the transform (each LHS and RHS) contained fewer atoms than the common core and only up to two cut locations (Table 5). This step was taken to to restrict the size, number, and location of the transforms to chemically meaningful replacements that are practically feasible. Transforms with at least three repeats were retained for further analysis.

## Acknowledgements

D.G. and U.Z. were supported by funding from the MRC (iCASE award MR/R015791/1 together with Helperby Ltd.). We thank Ian Gilbert, Ulrich Kleinekathöfer, Fabio Zuccotto, and Johannes Zuegg for insightful discussions.

## Author Contributions

DG and UZ conceived and designed the study, DG performed research, DG analysed and interpreted the data, DG wrote the manuscript, UZ edited the manuscript and supervised the research. Both authors read and approved the final manuscript.

## Data Availability

All the underlying code, together with all key datasets and working examples of generating results, can be accessed via Github: https://github.com/domgurvic/efflux_evaders_and_substrates

## Notes

### Competing Interest Statement

The authors have declared no competing interest.

