## Supplemental information for "Multidrug Efflux in Gram-Negative Bacteria: Rationally Modifying Compounds to Avoid Efflux Pumps"

### Supporting Information for: Multidrug Efflux in Gram-Negative Bacteria: Rationally Modifying Compounds to Avoid Efflux Pumps

Dominik Gurvic*,† and Ulrich Zachariae*,†,‡

†*Computational Biology, and*

‡*Biochemistry and Drug Discovery, School of Life Sciences, University of Dundee, Dundee, DD1 5EH, United Kingdom*

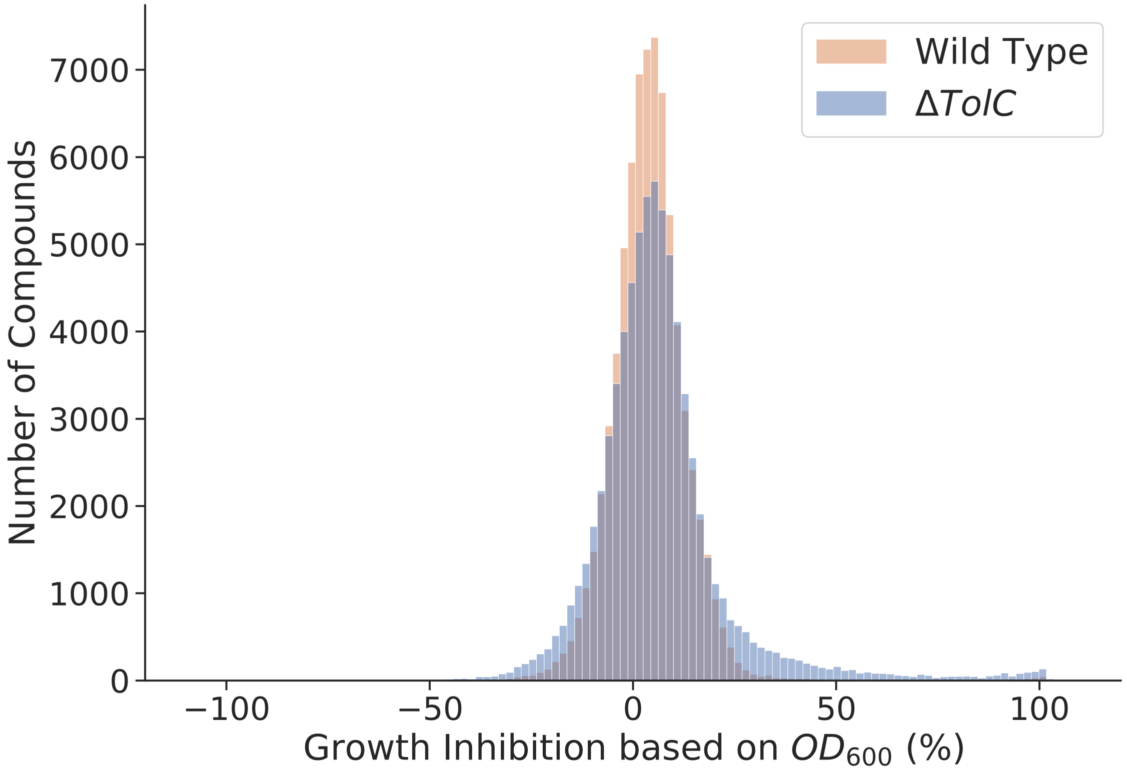


**Figure S 1.** Distribution of the growth inhibition activity of $\sim 74k$ compounds from the CO-ADD database tested against WT (orange) and efflux-deficient *tolC E. coli* (blue). The distribution for *tolC* data exhibits a lower peak and a distinct tail stretching to the right. This suggests that a number of compounds show greater activity in the efflux-deficient strain. A paired t-test between two distributions returned a p-value below 0.05, confirming that removing efflux pumps significantly affected growth inhibition.

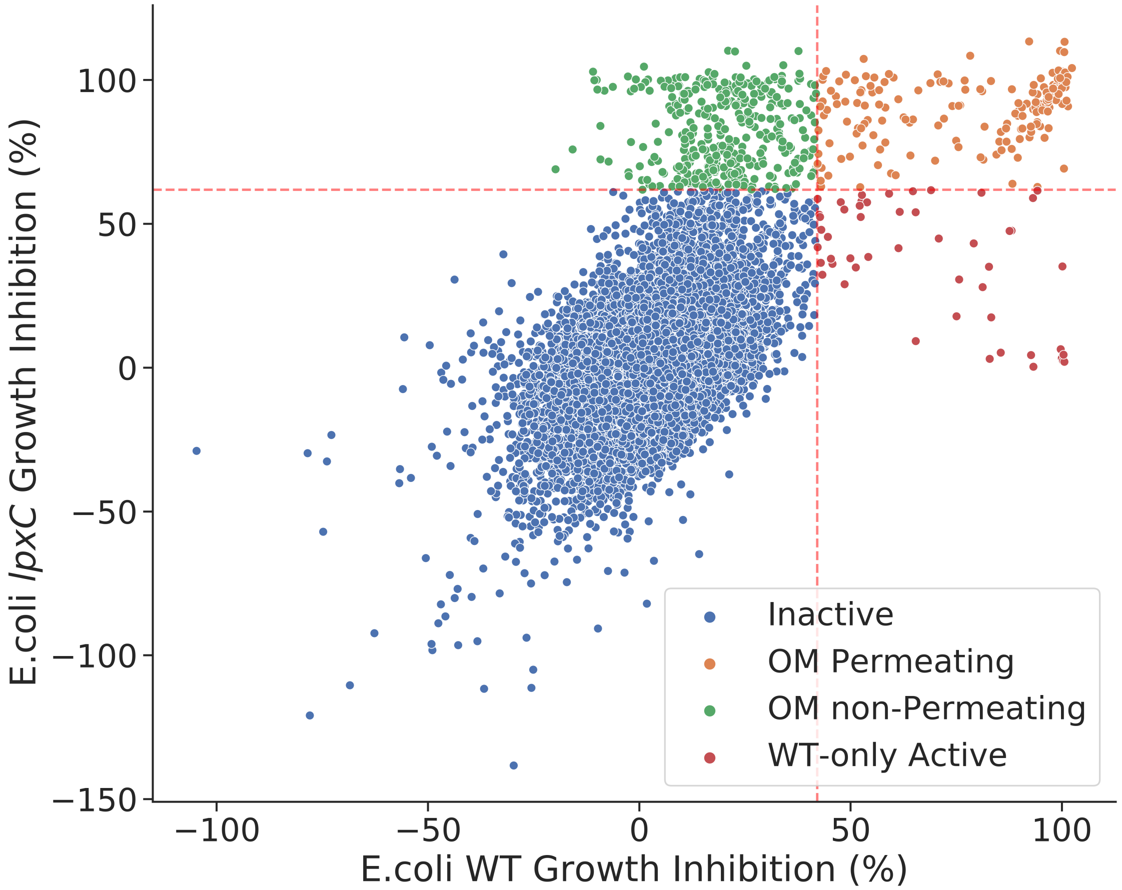


**Figure S 2.** We used the same activity thresholds as for WT vs. *tolC E. coli* to analyse GI activity in the WT and hyper-permeable *lpxC E. coli* strains. Inactives (blue) are identified as compounds that show activity below the threshold in both WT and *lpxC* strains. OM permeating compounds (orange) are active above the threshold in both WT and *lpxC*. OM non-Permeating compounds (green) are inactive in WT but active in *lpxC*. WT-only active compounds (red) are active only in the WT. The broken red lines show the activity thresholds at $GI > \mu + 4\sigma$ for both strains.

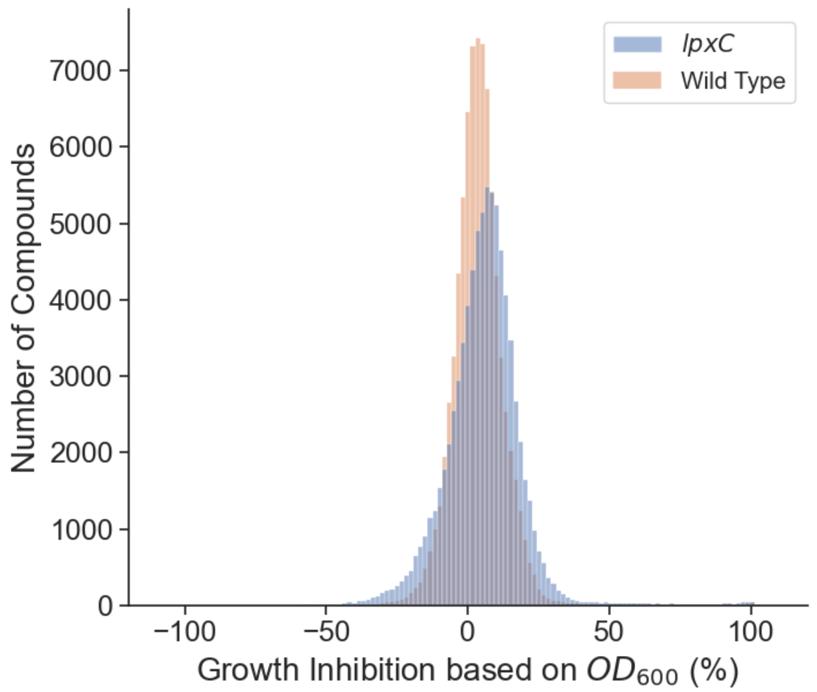


**Figure S 3.** Distribution of the growth inhibition activity of $\sim 74k$ compounds from the CO-ADD database tested against WT (orange) and hyper-permeable *lpxC E. coli* (blue). The mean of the distribution for *lpxC* data is clearly right-shifted, suggesting that, on average, the compounds show greater activity in the hyper-permeable strain. A paired t-test between two distributions returned a p-value below 0.05, confirming that OM permeability significantly affected growth inhibition.

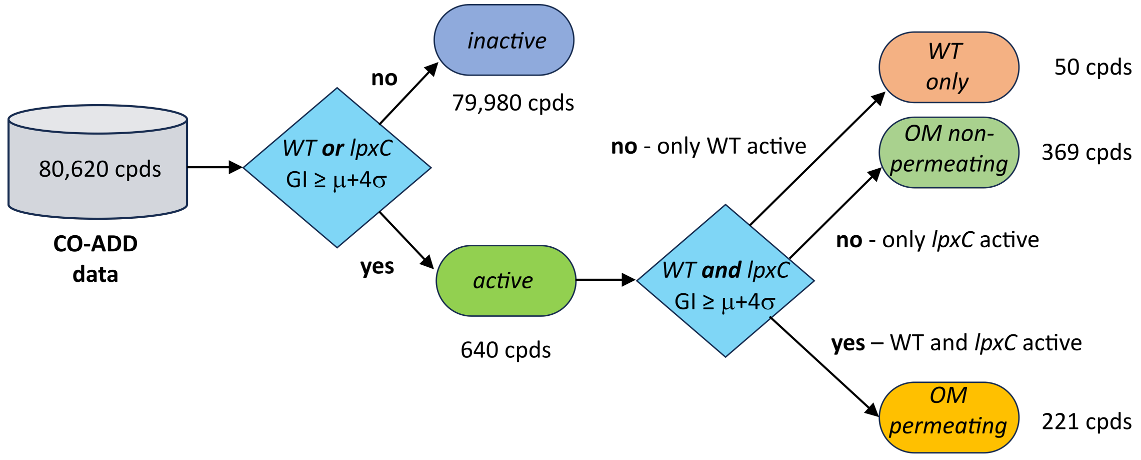


**Figure S 4.** Flow diagram of the classification scheme for OM permeable and non-permeable compounds, in addition to inactive and WT-only active compounds.

**Table S 1.** Average change in physicochemical features of OM non-permeable vs. permeable compounds according to WT and *lpxC E. coli* GI data.

| PC feature | OM non-permeating | OM permeating | Change |
|---|---|---|---|
| MW | 388.72 | 389.42 | +0.18% |
| *logP* | 2.87 | 2.49 | -13.24% |
| Rot. bonds | 4.88 | 4.45 | -8.81% |
| TPSA | 98.44 | 100.00 | +1.59% |
| HBA | 5.67 | 5.99 | +5.64% |
| HBD | 1.77 | 1.70 | -3.96% |

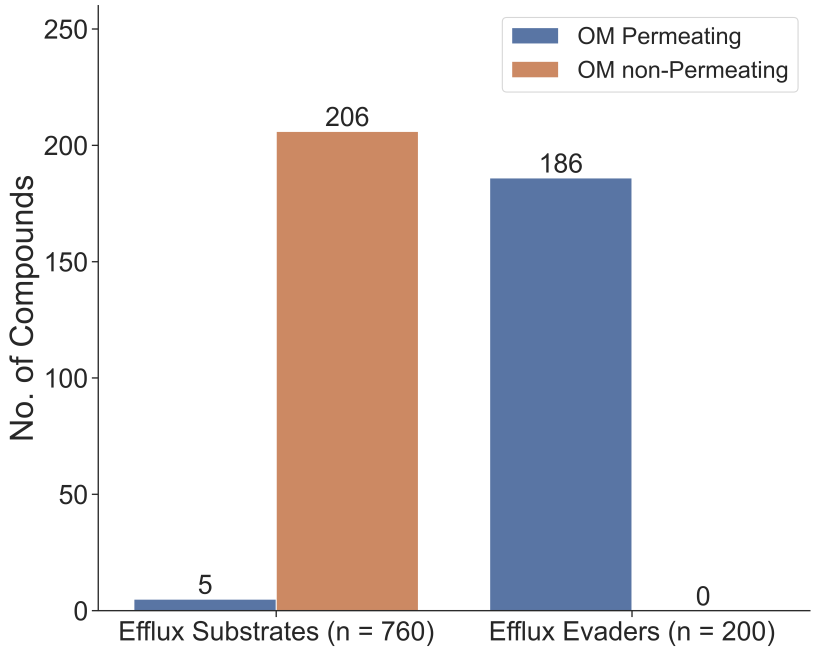


**Figure S 5.** Additional effect of OM permeation on efflux evaders and substrates. Out of 760 efflux substrates, 206 compounds are also classed as OM non-permeable, which means their WT inactivity could be caused by a combination of these features. Out of 200 efflux evaders, 186 are also classed as OM permeable.

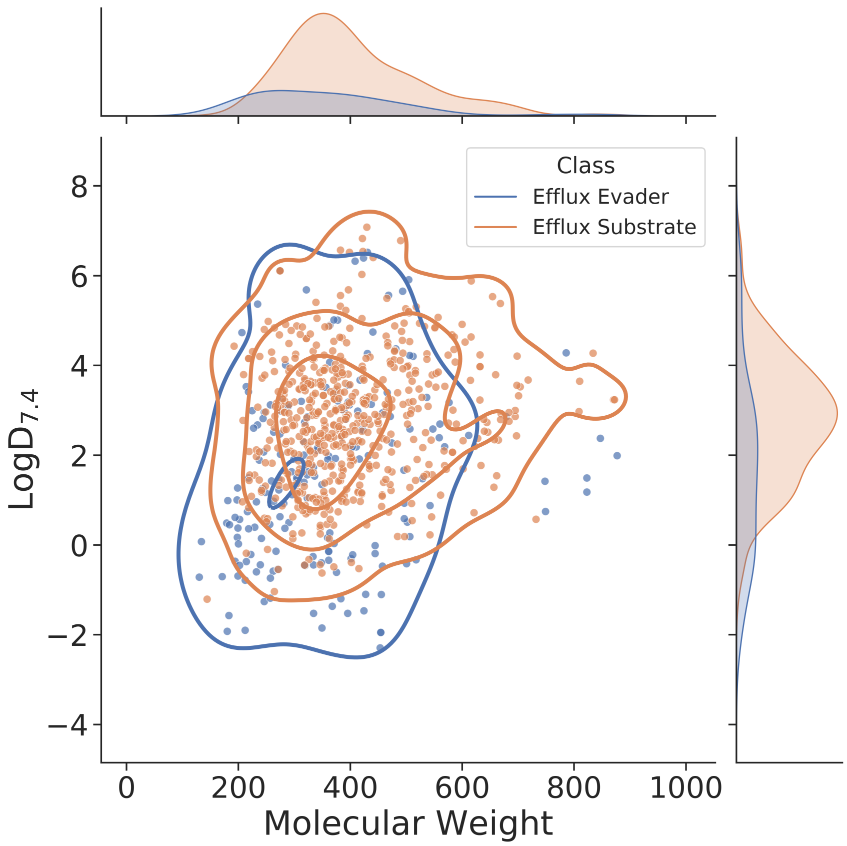


**Figure S 6.** Comparison of *LogD* and MW among efflux evaders and substrates show a shift to lower *LogD* and a slight shift to lower MW for efflux evaders.

| Cluster | Classes | MCS Evader | MCS Substrate | MCS All |
|---|---|---|---|---|
| 1 | Efflux Substrate - 37<br>Efflux Evader - 3 | 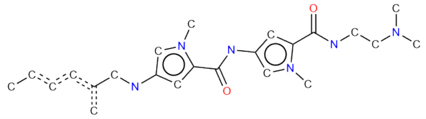 | 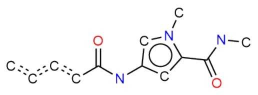 | 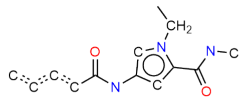 |
| 2 | Efflux Substrate - 9<br>Efflux Evader - 23 | 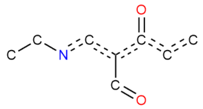 | 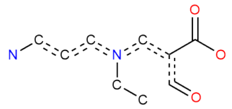 | 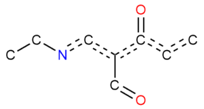 |

**Figure S 7.** Maximum common substructures (MCSs) of two distinct clusters of similar compounds resulting from t-SNE
